# Exploring groundwater microbial communities for natural attenuation potential of micropollutants

**DOI:** 10.1101/850750

**Authors:** Andrea Aldas-Vargas, Ernestina Hauptfeld, Gerben D.A. Hermes, Siavash Atashgahi, Hauke Smidt, Huub H.M. Rijnaarts, Nora B. Sutton

## Abstract

Groundwater is a key water resource, with 45.7% of all drinking water globally being extracted from groundwater. Maintaining good groundwater quality is thus crucial to secure drinking water. Micropollutants, such as pesticides, threaten groundwater quality which can be mitigated by biodegradation. Hence, exploring microbial communities in aquifers used for drinking water production is essential for understanding micropollutants biodegradation capacity. This study aimed at understanding the interaction between groundwater geochemistry, pesticide presence, and microbial communities in aquifers used for drinking water production. Two groundwater monitoring wells located in the northeast of The Netherlands and at 500 m distance from each other were sampled in 2014, 2015, 2016 and 2018. In both wells, water was extracted from five discrete depths ranging from 13 to 54 m and used to analyze geochemical parameters, pesticide concentrations and microbial community dynamics using 16S rRNA gene sequencing and qPCR. Groundwater geochemistry was stable throughout the study period and pesticides were heterogeneously distributed at low concentrations (μg/L range). Integration of the groundwater chemical and microbial data showed that geochemical parameters and pesticides exerted selective pressure on microbial communities. Furthermore, microbial communities in both wells showed a more similar composition in the deeper part of the aquifer as compared to shallow sections, suggesting vertical differences in hydrological connection. This study provides initial insights into microbial community composition and distribution in groundwater systems in relation to geochemical parameters. This information can contribute for the implementation of bioremediation technologies that guarantee safe drinking water production from clean aquifers.

**Importance section:** Groundwater is an essential source of drinking water. However, its quality is threathened by the presence of micropollutants. Certain microorganisms are capable of degrading micropollutants. However, groundwater is an unexplored environment, where the biodegradation potential of naturally-present microorganisms is unknown. We thus explore how groundwater microbial ecology in shaped by groundwater composition, namely geochemical parameters and micropollutants. This is a first step towards understanding which microbial communities and environmental conditions support natural attenuation of micropollutants. This study thus provides a first step towards developing in situ bioremediation strategies to remove micropollutants from groundwater used for drinking water production.

## 1. Introduction

Groundwater is a key water resource, with 45.7% of all drinking water being abstracted from groundwater globally (1). Therefore, maintaining good groundwater quality is crucial to maintaining drinking water security. One group of micropollutants (MP) that threatens groundwater quality is formed by pesticides. In 2016, more than 4 billion kilograms of pesticides were used for agricultural purposes worldwide (2). Pesticides seep into groundwater, resulting in concentrations that often exceed regional legislations for drinking water production (3). Post extraction treatment by advanced water treatment technologies, such as adsorption to activated carbon or advanced oxidation can effectively remove pesticides in some cases, but are neither cost-effective nor suitable for all situations (4).

As an alternative, in situ biodegradation or natural attenuation of MP, mediated by indigenous microorganisms, is regarded as a more sustainable means for safeguarding drinking water resource quality (5). Harnessing the biodegradation capacity of aquifers however, is especially challenging, due to oligotrophic conditions, due to low DOC concentrations, and limited oxygen availability. Oxygen, as the most favorable electron acceptor, is often quickly depleted below the surface, especially in relatively young geological formations as found in river flood plains and delta’s; in these deeper aquifers, microbes rely on other available electron acceptors like nitrate, Fe (III), sulfate, etc. Groundwater is also typically depleted in nutrients and assimilable organic carbon, because preferable carbon substrates are consumed by microbes in mineral soil and the vadose zone (6). A potential degrader therefore has to compete with other microbes for scarce assimilable organic carbon for energy, while still maintaining efficient degradation activity (3). Whereas biodegradation of pesticides has been shown in soil, activated sludge, and aerobic sand filter systems (7–12), few studies have been conducted in low-concentration, low-biomass environments that could be representative for groundwater (3). Harnessing the biodegradation capacity of aquifers, i.e. for PM removal, necessitates a thorough biogeochemical understanding of these oxygen-limited and oligotrophic ecosystems.

Recent studies have tried to describe microbial communities in groundwater ecosystems in situ, or in microcosm experiments that simulated environmental conditions (13–20). Results from high-throughput sequencing of 16S rRNA gene amplicons show that a large number of microorganisms in groundwaters cannot be classified even on class or phylum level (13, 17), which largely precludes assumptions about their biodegradation potential. Whereas some studies examined the correlations among microbial community, pollutant abundance, and geochemical parameters (13, 18–20), these datasets did not consider stabilities in microbial community and redox conditions within the aquifer, over time and spatially. Considering the diversity of electron acceptors, dissolved organic carbon (DOC) as electron donor, and low nutrient concentrations in groundwater, long-term repeated sampling of microbial communities in a variety of groundwater compositions at a fixed set of locations and depths is required to make conclusions about MP natural attenuation distributions over aquifers. To this end, this study aimed to elucidate the interaction of groundwater geochemistry, pesticide presence, and aquifer microbial communities to understand the potential for the natural attenuation of pesticides. Geochemical properties and pesticide concentrations were measured from groundwater monitoring wells in the northeast of The Netherlands over the course of several years, and 16S rRNA gene amplicon sequencing was used for microbial community analysis, to understand aquifer biogeochemistry. Each well was sampled at five different depths between 12m and approximately 60m below the surface. The results presented here provide insight into microbial community composition and how it relates to groundwater composition, with the ultimate aim of better understanding the aquifers’ potential for natural attenuation.

## 2. Materials and Methods

### 2.1 Site description

The groundwater monitoring wells used for this study are located in an agricultural area in the northeast of The Netherlands. Monitoring wells are located around groundwater extraction locations and used for drinking water production. To ensure the water quality of drinking water, several monitoring wells are installed in the surroundings of extraction locations. Two of those monitoring wells were sampled for this study. A map of the wells’ locations is provided in Supplementary Information (Figure S1). The first well (designated as well 22) is further upstream form the extraction location and adjacent to a canal, while the second one (designated as well 23) is closer to the extraction location. The distance between well 22 and 23 is around 500 m, and both wells are filtered at five discrete depths from 13-54 m below ground level (Table S1).

### 2.2 Groundwater sampling

Groundwater sampling was performed at the above-mentioned monitoring wells at each discrete depth. Samples were collected per depth in duplicates for the years 2014, 2015, 2016 and in singletons for 2018. Before sample collection, wells were flushed by extraction, and the volume of water contained per pipe was discarded three times before sampling. Turbidity was measured during this process, and the samples were taken after the turbidity measurement stabilized below 1 NTU. Samples were collected in 10 L jerry cans and stored at 4°C before further analysis.

### 2.3 Geochemical and micropollutant quantification

Geochemical parameters and MP had been monitored starting in 2000 by an accredited Dutch Government Laboratory. Samples for these analyses were taken by the same laboratory according to specific requirements of the different analytical techniques. The DOC was determined by combustion in accordance with NEN-EN 1484:1997 (21). The iron (II) concentration was analyzed using an inductively coupled plasma and mass spectrometry (ICP-MS) in accordance with NEN-EN-ISO 17294-2: 2004 (22).The ammonium (N) concentration was determined with a discrete analyzer in accordance with NEN-ISO 15923-1: 2013 (23) and the nitrate (N) concentration by ion chromatography (IC) (24). The sulfate concentration was determined in accordance with NEN-EN-ISO 10304-1:1995 (25). The pesticides bentazon, mecoprop, chloridazon and the metabolites chloridazon-desphenyl (CLZD) and chloridazon-methyl-desphenyl (CLZMD) were quantified by liquid chromatography coupled with a mass spectrometer (LC-MS). After acidification and addition of labeled internal standards, the samples were injected and analyzed (24). 1,4-dioxane (henceforth dioxane) was determined by using purge and trap gas chromatography coupled with a mass spectrometer (GC-MS), and the metabolite 2,6-dichlorobenzamide (henceforth BAM) was measured in a GC-MS triple quad (QQQ) (24).

### 2.4 Groundwater filtration

In order to concentrate biomass, groundwater samples were filtered through Isopore membrane filters with a pore size of 0.2 μm (Merck Group, Darmstadt, Germany). The exact volume of water filtered per sample is given in Tables S2 and S3. Filters were snap frozen in liquid nitrogen and stored at −20°C prior to DNA extraction. The glassware from the filtration equipment was cleaned with absolute ethanol between each filtration process.

### 2.5 DNA extraction and quantification, PCR amplification and sequencing of the 16s rRNA gene, and quantitative PCR (qPCR)

Microbial DNA was extracted from each filter using the DNeasy PowerSoil Kit (Qiagen, Hilden, Germany) according to the manufacturer’s instructions. The quality of DNA (average molecular size) was controlled on 1% (w/v) agarose gels stained with 1× SYBR^®^ Safe ethidium bromide (Invitrogen, Grand Island, NY) and its quantity was measured using the dsDNA HS Assay kit for Qubit fluorometer (Invitrogen).

The PCR amplification for samples from 2014 to 2016 was conducted with primers 27F (GTTYGATYMTGGCTCAG) and 338R (GCWGCCWCCCGTAGGWGT) targeting the V1 and V2 regions of the 16S rRNA gene. However, primers targeting regions V1-V2 have recently been shown to be better suited for gut microbiota studies rather than for environmental samples (26). Therefore, for the 2018 dataset, the primers used were 515F (GTGCCAGCMGCCGCGGTAA) and 806R (GGACTACHVGGGTWTCTAAT) targeting the V4 region (27).

#### 2.5.1 Set of samples 2014-2016

In this study, a two-step PCR protocol was used. With this approach, tags and adapters were added in a second round of PCR amplification (Tables S4 and S5).

The initial PCR mix was prepared as described in Table S6 and the second PCR mix as in Table S7. The amplification program for both PCR mixes is detailed in Table S8. The quality and concentration of the PCR products were determined through analysis on a 1% (w/v) agarose gels stained with 1×SYBR^®^ Safe ethidium bromide (Invitrogen) and using the dsDNA HS Assay kit for Qubit fluorometer (Invitrogen). The barcoded samples were pooled in equimolar concentrations and sent for sequencing on an Illumina Miseq machine (GATC-Biotech, Konstanz, Germany; now part of Eurofins Genomics Germany GmbH). Sequence data were submitted to the European Bioinformatics Institute under study accession No PRJEB34986. The barcode sequence for each sample is detailed in Table S9.

qPCR analysis was used to quantify total bacteria and archaea based on the 16S rRNA gene, and functional genes involved in nitrate reduction (*nirS*, *nirK*, *nosZ*) (28) and sulfate reduction (*dsrB*) (29). Analysis was performed on an iQ SYBR Green using Bio-Rad super mix using CFX384 Touch™ Real-Time PCR Detection System. All qPCR assays were performed in triplicate with a total volume of 10 μL reactions. Gene copy numbers were calculated per ml groundwater. Detailed information of the qPCR primers and amplification protocols can be found in Table S10.

#### 2.5.2 Set of samples 2018

The PCR mix was prepared as described in Table S11, and the PCR program used is detailed in Table S12. The PCR products were cleaned with the MagBio Beads Cleanup Kit (MagBio, MD, USA) according to the manufacturer’s instructions. Quality and concentration of the PCR products were determined as described in Section 2.5.1. Samples were pooled (total of 72) in approximately equal concentrations (4 ×10^6^ copies μl^−1^) to ensure equal representation of each sample. The barcode sequence for each sample is detailed in Table S13. The pooled samples were cleaned with the MagBio Beads Cleanup Kit (MagBio) for a second time and quantified again using the dsDNA HS Assay kit for Qubit fluorometer (Invitrogen). The resulting library was sent to Eurofins genomics (Konstanz, Germany) for 2X150nt sequencing on an Illumina Hiseq2500 instrument. These sequence data were submitted to the European Bioinformatics Institute under study accession No XXXXX and sample accession No XXXXX.

### 2.6 Data processing and analysis

Sequence analysis of the raw data was performed in NG-Tax using default settings (30). In short: paired-end libraries were demultiplexed using read pairs with perfectly matching barcodes. Amplicon sequence variants (ASV) were picked as follows: for each sample sequences were ordered by abundance and a sequence was considered valid when its cumulative abundance was ≥ 0.1%. Taxonomy was assigned using the SILVA reference database version 128 (31). ASVs are defined as individual sequence variants rather than a cluster of sequence variants with a shared similarity above a pre-specified threshold such as operational taxonomic units (OTUs). Even though the read length of the Miseq and Hiseq data differed (250nt vs 150nt respectively) NG-Tax analyzed the same read lengths (140nt) for both datasets to homogenize the data.

Analysis of microbial communities was performed in R version 3.5 (32). Alpha diversity (within sample diversity) metrics were calculated using Faith’s phylogenetic diversity (33), implemented in the picante (34) package. Beta-diversity (between sample diversity) was calculated using weighted and unweighted UniFrac in the phyloseq package (35). To determine the univariate effects of the geochemical environment and MP on the microbial composition, redundancy analysis (RDA) was performed using the rda functions from vegan (36). This is a gradient analysis technique which summarizes linear relationships between multiple components of response variables (microbes) explained by a set of explanatory variables (geochemical components and pollutants) by multiple linear regression of multiple response variables on multiple explanatory variables. To determine which of the electron acceptors and MP significantly explained the most variance in the composition of microbial communities and which variables generated the most parsimonious model, forward and backward automatic stepwise model selection were performed using the ordistep function. The same automatic model selection was performed with the ordiR2step function (only forward selection and different weight for R2adj and p-values) to ensure a robust model. To determine the shared variation explained by the electron acceptors and selected MP, variation partitioning was performed using the varpart function from vegan.

## 3. Results

### 3.1 Groundwater Geochemical Composition

Per sample location and depth, groundwater geochemical parameters, such as DOC, iron (II), ammonium, nitrate and sulfate, remained stable over the course of 16 years (Figure 1, a and b). All filters in the sampled aquifers qualify as hypoxic with oxygen levels below 0.5 mg L^−1^ at almost all sampling times (Supplementary Table S14). Furthermore, clear zonation of the availability of different electron acceptors was observed. For example, in well 23, nitrate was present at filters 1-3 (13-37m), at average concentrations of 18, 15, and 12 mg L^−1^, respectively (Figure 1b). A low concentration of iron (II) was observed, suggesting that nitrate is most likely the dominating electron acceptor. This was further supported by the qPCR data (Figure 1d), which shows a higher abundance for the genes *nirS* and *norZ*, involved in nitrite and nitrous oxide reduction, respectively, in filters 1-3 of well 23 compared to the other samples. In filters 4 and 5 of well 23 (47 and 54m), iron (II) was observed at average concentrations of 11.4 and 14 mg L^−1^, indicating iron-reduction is occurring in the deeper zones. This was accompanied by a 1-2 order of magnitude drop in *nirS* and *nosZ* abundance compared to filters 1-3 of well 23, suggesting that nitrate is not used as a primary electron acceptor. In contrast, well 22 was characterized by more anoxic conditions, with no nitrate measured during 16 years of monitoring. Furthermore, well 22 had less clear zonation of electron acceptors (Figure 1a). Both iron (II) and sulfate were present and their concentrations varied between filters (iron (II) 0.4-14.7 mg L^−1^, sulfate 0-93 mg L^−1^), most likely indicating a mix of iron- and sulfate-reducing conditions. Sulfate-reduction appeared to be most active in filters 22-1 and 22-2 as indicated by the low concentration of sulfate compared to filters 22-4 and 22-5. A high abundance of the *dsrB* gene (Figure 1c), which is involved in sulfate reduction, provided further evidence for sulfate as an electron acceptor in filters 22-1 and 22-2.

**Figure 1.**
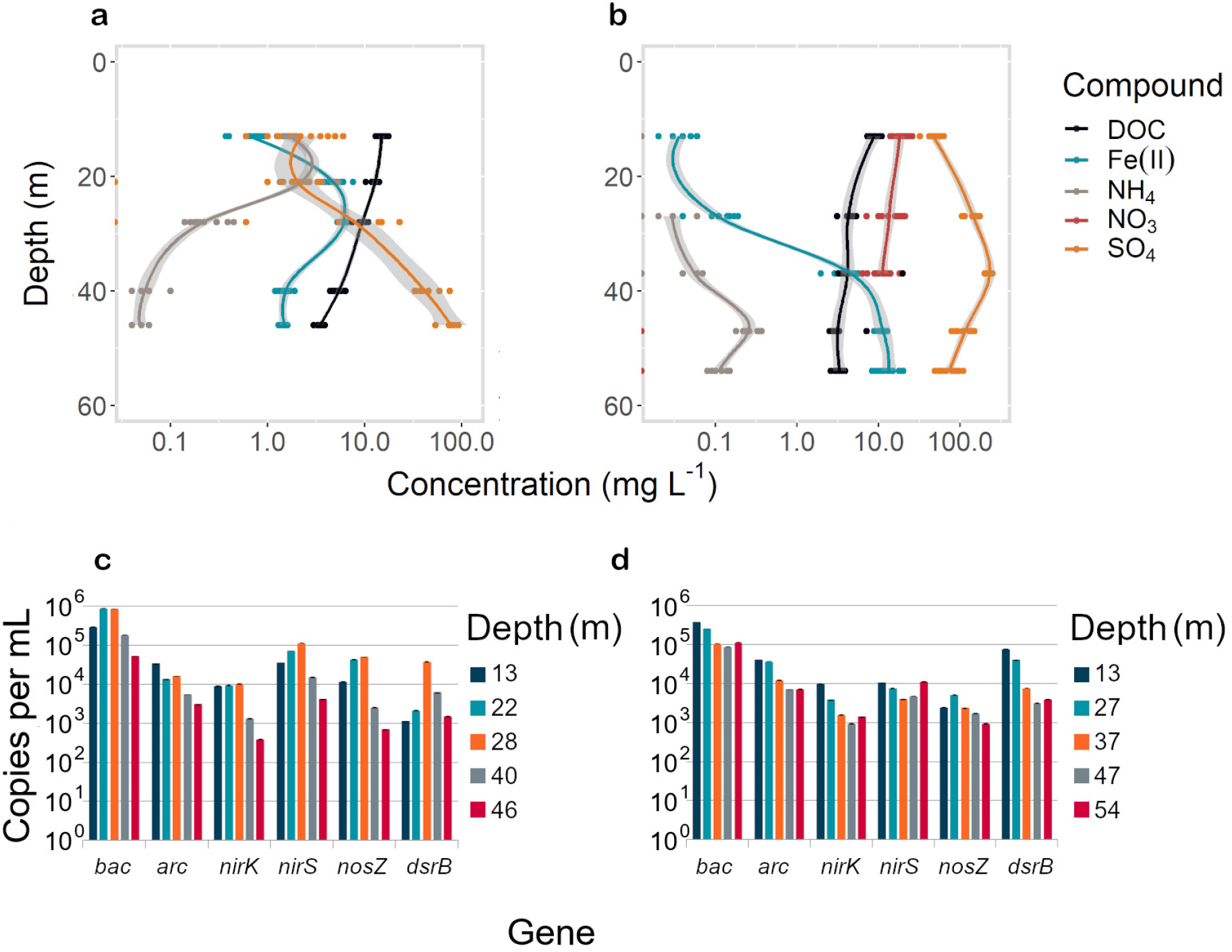
Top: Geochemistry of wells 22 (a) and 23 (b). Each point of the same color represents one measurement in one year between 2000 and 2016. The lines represent the average of all measurements. The grey area on both sides of the lines represent the loess confidence intervals. measurements were taken at least one year apart. Bottom: qPCR measurements of bacteria (eub), archaea (arc) 16S rRNA genes and functional genes of denitrification (*niK, nirS*, and *nosZ*) and sulfate reduction (*dsrB*) in wells 22 (c) and 23 (d).

The groundwater geochemical data showed zonation in electron acceptor availability coupled with DOC as the dominating electron donor. DOC was found in both well 22 and 23 at concentrations between 2.4 and 19.7 mgL-1 (Figure 1, a and b). In both wells, DOC concentrations decreased with filter depth, with the largest decrease observed in well 22.

Regular monitoring of hundreds of organic MP by the drinking water company (full data not shown) revealed that seven compounds were consistently encountered in wells 22 and 23. However, in contrast to the stability of geochemical components, the concentrations of the MPs varied greatly within the same filters as well as between filters (Figure 2). Three pesticides (bentazon, chloridazon, mecropop), three pesticide transformation products (BAM, CLZD, CLZMD), and one solvent (dioxane) exceeded the threshold (0.1 μg L^−1^) of the European framework for groundwater quality (37) on at least one occasion. We found concentrations ranging from below the detection limit to 8.4 μg L^−1^. All measured pollutants were detected in both wells in at least one measurement except chloridazon, which was only detected in well 22. The most abundant MP was dioxane with an average of 0.2μg L^−1^ in well 22 and 1.0μg L^−1^ in well 23. CLZD, a stable metabolite of the pesticide chloridazon, was particularly abundant in well 22 and observed at concentrations of between 0.3 and 1.6 μg L^−1^.

**Figure 2.**
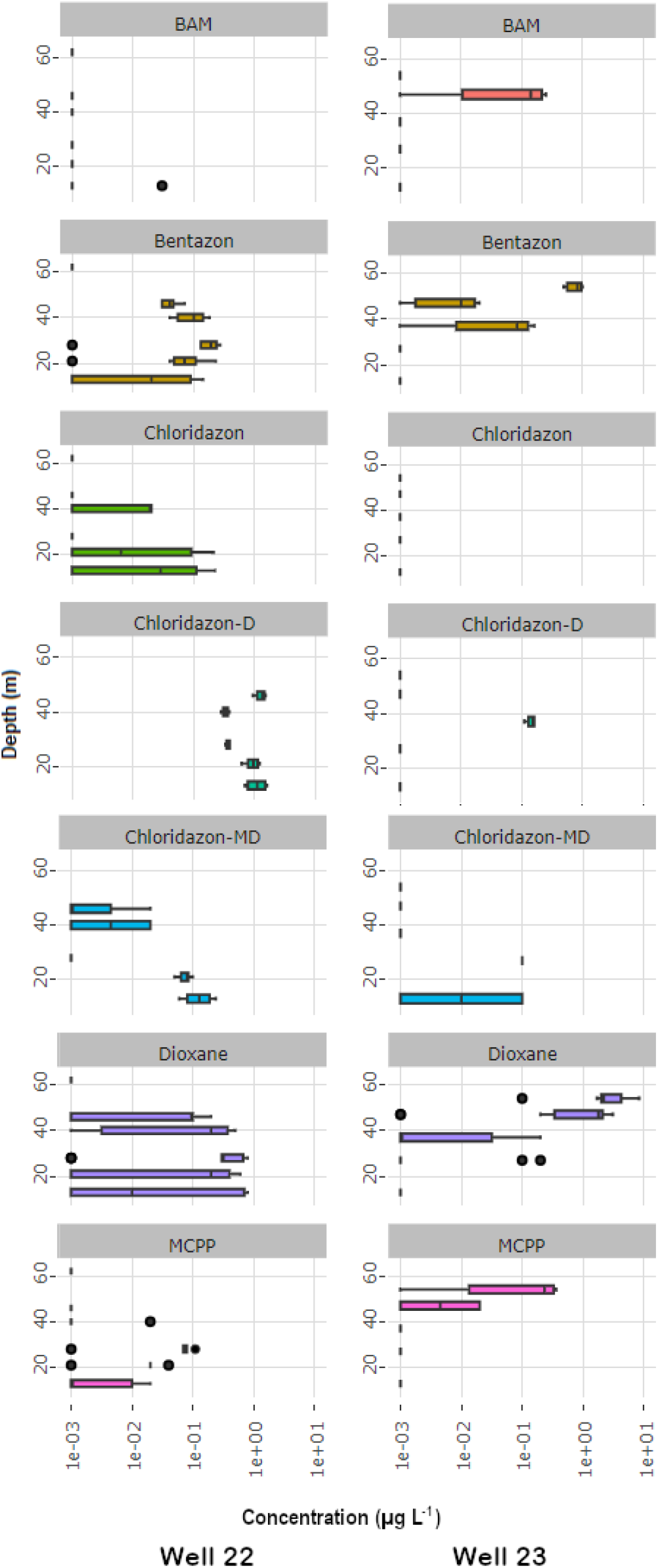
concentrations of micropollutants in wells 22 and 23 from 2000 to 2016. The box plots represent series of 4-16 measurements. The measurements were taken at least one year apart.

### 3.2 Microbial community composition

OTU-classification was performed using the NG-Tax pipeline (30). A large portion (7-55%) of OTUs could not be classified even at phylum level (Figure 3a). No OTUs were assigned to the domain of *Archaea*. On average, the most abundant phyla in well 22 were *Proteobacteria* (26.8±14.9%), *Chloroflexi* (11.8±5.7%), *Candidatus Omnitrophica* (8.0±3.2%), *Nitrospirae* (7.5±12.1%), *Bacteroidetes* (4.0±3.3%), *Firmicutes* (3.3±1.5%), *Microgenomates* (2.2±2.2%), *Nitrospinae* (2.4±3.8%), and *Parcubacteria* (1.7±1.3%, Figure 3a). The most abundant phyla of well 23 were classified as *Proteobacteria* (25.6±8.2%), *Omnitrophica* (16.9±9.2%), *Microgenomates* (8.4±4.0%), *Nitrospirae* (7.9±9.2%), *Chloroflexi* (6.6±4.8%), *Ignavibacteriae* (4.6±6.6%), *Parcubacteria* (1.5±1.4%), *Bacteroidetes* (1.1±1.3%), and *Acidobacteria* (0.9±1.5%, Figure 3b).

**Figure 3.**
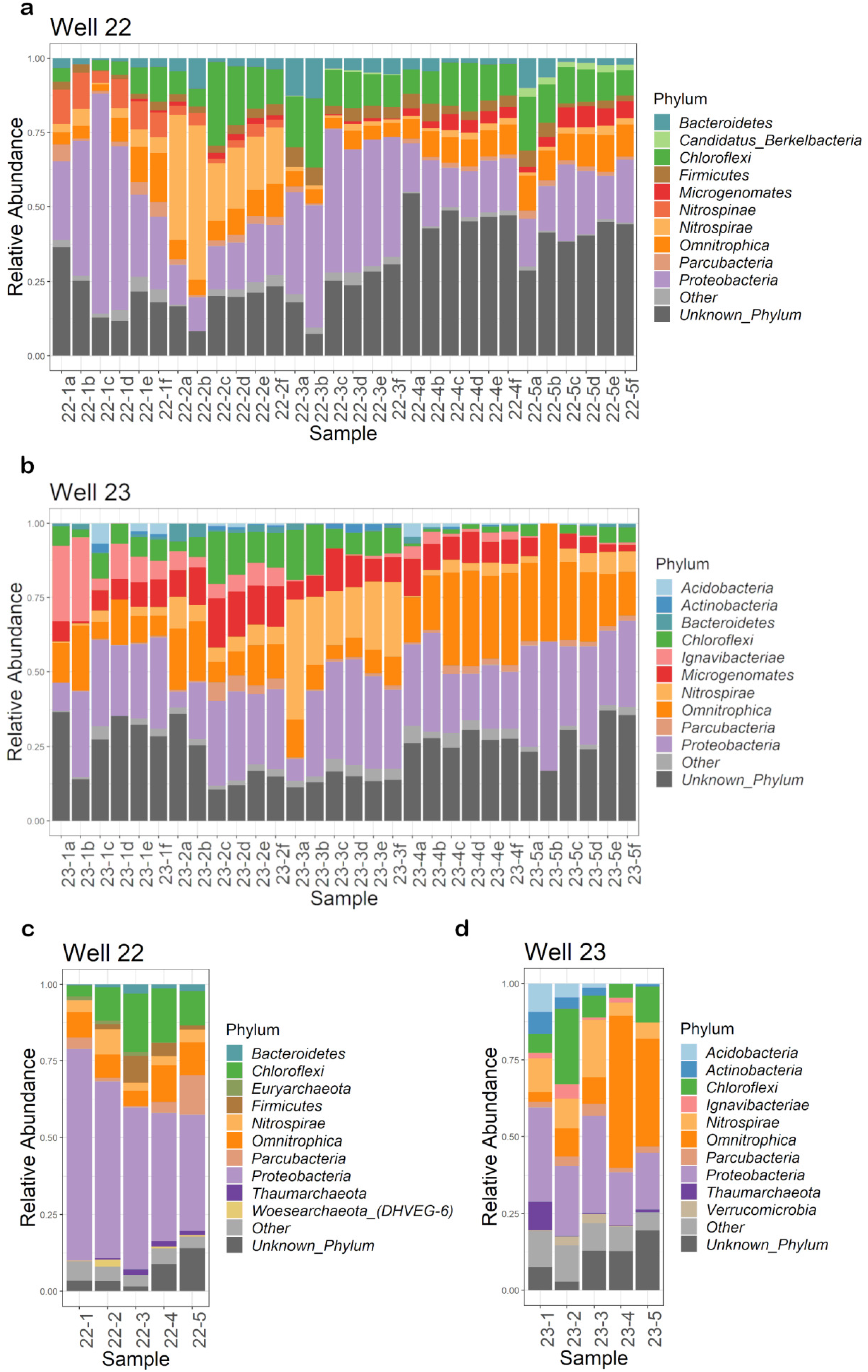
Relative abundance of the top 10 microbial phyla of well 22 (a, c) and well 23 (b, d) from the samples collected in 2014-2016 (a & b) and 2018 (c & d). “Unknown_Phylum” stands for all sequences that could not be classified at phylum level. “Other” are all the phyla that are classified but do not belong to the top 10.

For the samples taken in 2018, the portion of sequences that could not be assigned to any phylum ranged from 1.4-19.4%. The most abundant phyla in well 22 were *Proteobacteria* (51.7±12.5%), followed by *Chloroflexi* (12.6±6.2%), *Candidatus Omnitrophica* (8.8±2.8%), *Nitrospirae* (4.3±2.2%), *Parcubacteria* (4.3±5.0%), *Firmicutes* (3.2±3.5%), *Bacteroidetes* (1.5±1.1%), *Thaumarchaeota* (1.1±0.8%), *Euryarchaeota* (0.7±0.6%), and *Woesearchaeota_(DHVEG-6)* (0.7±0.9%, Figure 3c). The most abundant phyla of well 23 were *Proteobacteria* (24.1±6.7%), *Omnitrophica* (21.1±20.2%), *Chloroflexi* (10.8±8.1%),*Nitrospirae* (9.8±5.8%), *Acidobacteria* (3.1±3.8%), *Actinobacteria* (2.9±2.8%), *Parcubacteria* (2.5±1.1%), *Thaumarchaeota* (2.2±3.9%), *Ignavibacteriae* (1.8±1.8%), and *Verrucomicrobia* (1.3±1.6%, Figure 3d).

### 3.3 Beta-diversity

For the samples from 2014-2016, average unweighted Unifrac distances were calculated between the six samples from each filter. Furthermore, the average unweighted Unifrac distances between the six samples from each filter and the six samples of any other filter were calculated (Figure 4). When displaying all average values in a heatmap, a clear diagonal was visible, indicating that the average dissimilarity within samples (n 6; duplicates from three consecutive sampling campaigns) from the same filters was the smallest for each of the ten measured filters (average Unifrac distance 0.38±0.18). Dissimilarity did not increase from samples taken one year apart to samples taken two years apart. This indicated that the microbial communities at each location and depth remained fairly stable over the course of the three measured years. Filter 23-4 appeared to have the most stable community within a filter with an average dissimilarity value of 0.30±0.08 while 23-5 was the least stable (0.53±0.27). Unifrac distances were smaller than average between filters 1, 2, and 3 of well 22 and filters 1, 2, and 3 for well 23, as well as between the filters 4 and 5 across wells 22 and 23. These results indicate that the microbial communities from the filters 4 and 5 were more similar across the two wells than the microbial communities from shallow and deep filters within a single well. These observations were further confirmed using principle coordinate analysis using unweighted Unifrac distances for all 16S rRNA gene sequence-based microbial profiles in samples taken from 2014-2016 shows a clear division of the 60 samples into three groups. The first cluster in the top left quarter of figure 5a is made up of samples of the two deepest filters (filters 4 and 5) of both wells 22 and 23. It should be noted that 23-5 had the largest Unifrac distance within the sample group, and that sample 23-5b appears to be an outlier. The second cluster in the bottom left quadrant of the figure is comprised of samples from the top three filters of well 22 (Figure 5a). The third cluster in the middle right of the figure consists of samples from the top three filters of well 23. To minimize possible primer bias, new samples were collected in 2018 and the analysis was repeated using new 16S rRNA primers targets the variable region V4 (Figure 5b). The results showed the same distribution of the samples as the analysis of samples taken in earlier years targeting variable regions V1-V2 of the 16S rRNA gene, with the samples from deep filters from both wells clustering towards the top left, the shallow samples from well 22 towards the bottom left, and the shallow samples from well 23 clustering in the middle right part of the figure.

**Figure 4.**
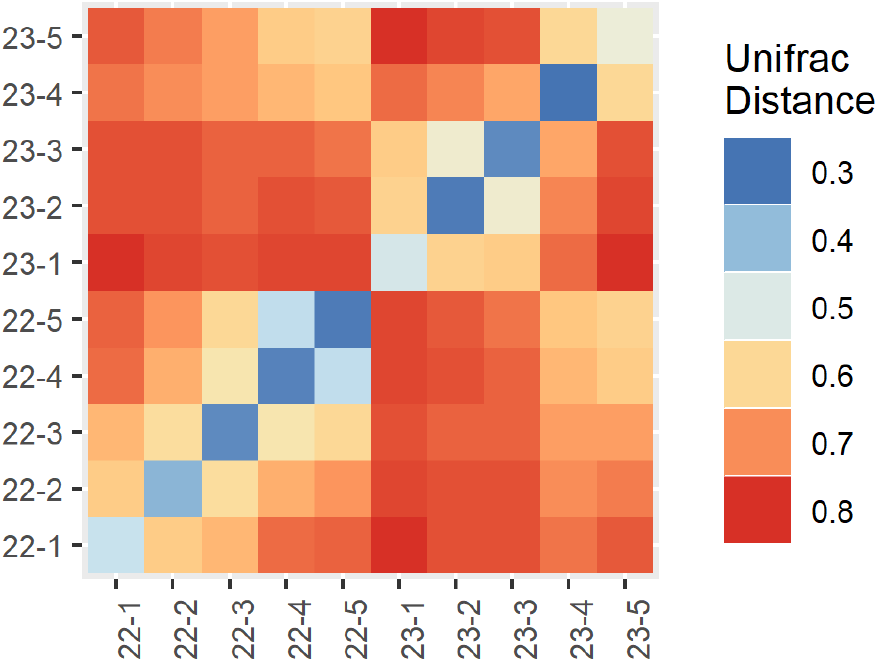
Heatmap of unweighted Unifrac distances. The filters are displayed on both the x and the y-axis. Filters with higher similarity (low distance) are displayed in light orange and blue, filters with lower similarity (large distance) are displayed in dark orange and red. To compute the average distances, we calculated the Unifrac distances between each sample of one filter compared to each sample of another filter. The arithmetic mean of all comparisons between two filters is displayed as Unifrac Distance in this figure.

**Figure 5.**
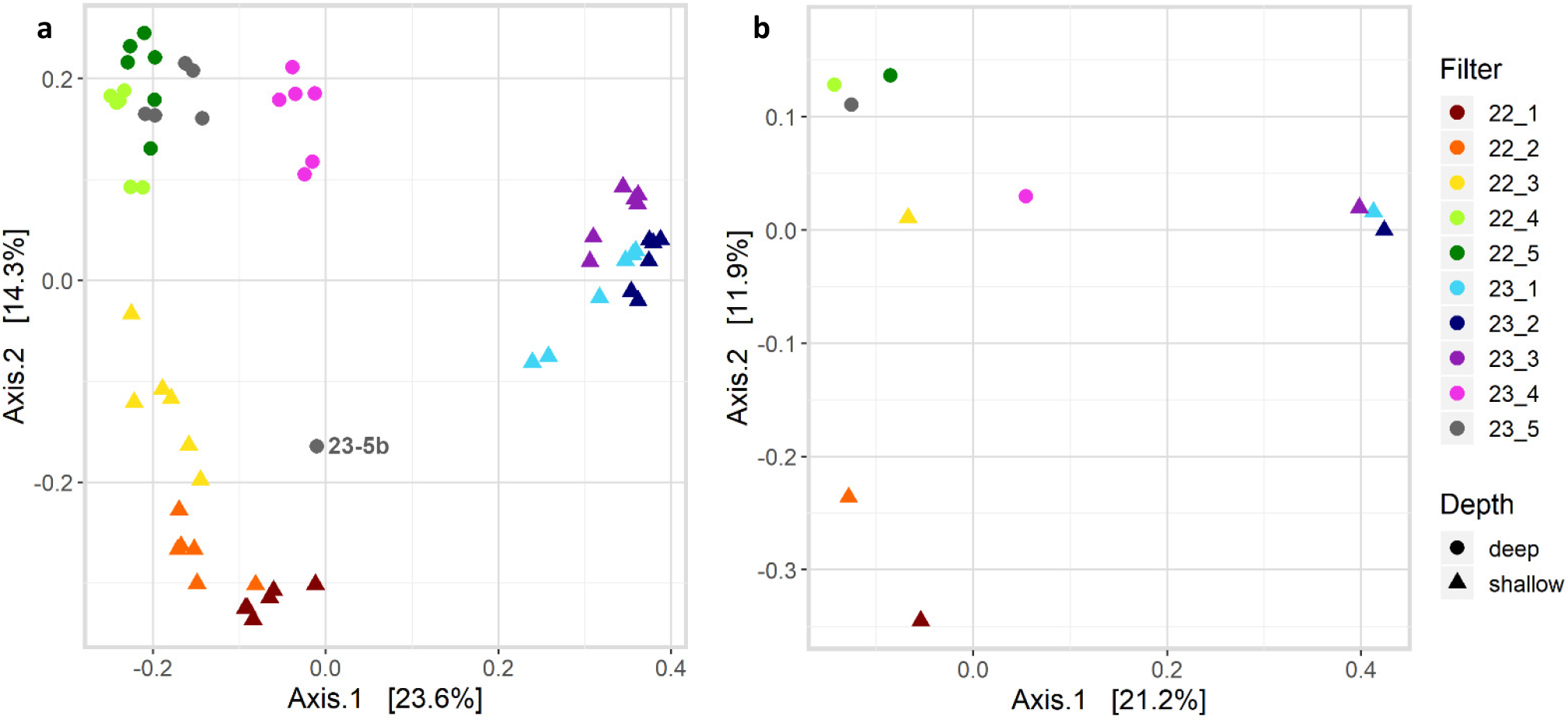
Principle coordinate analysis based on pairwise unweighted Unifrac distances of microbial profiles of samples taken 2014-2016, sequenced at region V1-V2 (a), and samples taken in 2018, sequenced at variable region V4 (b) of the 16S rRNA gene.

### 3.4 Alpha-diversity

Faith’s phylogenetic diversity and species richness of the sampled microbial communities was computed using R. The highest diversity was found in sample 23-5d (29.2 Phylogenetic Diversity, 235 ASVs detected). The sample with smallest phylogenetic diversity belonged to sample 23-5b (2.6) and contained only 5 (Supplementary Table S15). Microbial communities observed in filters 4 and 5 (22.8±5.4) of both wells were more diverse than those in filters 1-3 (18.5±4.7) (p=0.002 two-sided t-test).

### 3.5 Redundancy Analysis

To determine to what extent concentrations of geochemical components and pollutants on microbial communities can explain the observed variation in microbial composition, we used RDA. Out of all the measured components, the most variation was explained by nitrate, sulfate, iron (II), DOC, ammonium, dioxane, and CLZD. The resulting model explained 53% of the variance between the microbial communities (Figure 6a). All the predictors show high significance (p-value<0.001) in the model. Out of the variables in the model, the most variation is explained by nitrate, followed by DOC, iron (II), sulfate, dioxane, ammonium, and CLZD. The microbial group most associated with the nitrate was family FW13 from the phylum *Nitrospirae*. The family *Nitrospiraceae* of the phylum *Nitrospirae* closely correlated with the presence of ammonium. Two members of the phylum *Proteobacteria*, one belonging to the order *Methylococcales* and one affiliated with the genus *Synthrophus*, correlated with both iron (II) and ammonium concentration. The samples clustered in three groups: the samples from the shallow filters in well 23 are grouped over the arrow for nitrate; samples of the shallow filters in well 22 appear close to DOC; all samples from deep filters are located between sulfate, dioxane, and iron (II).

**Figure 6.**
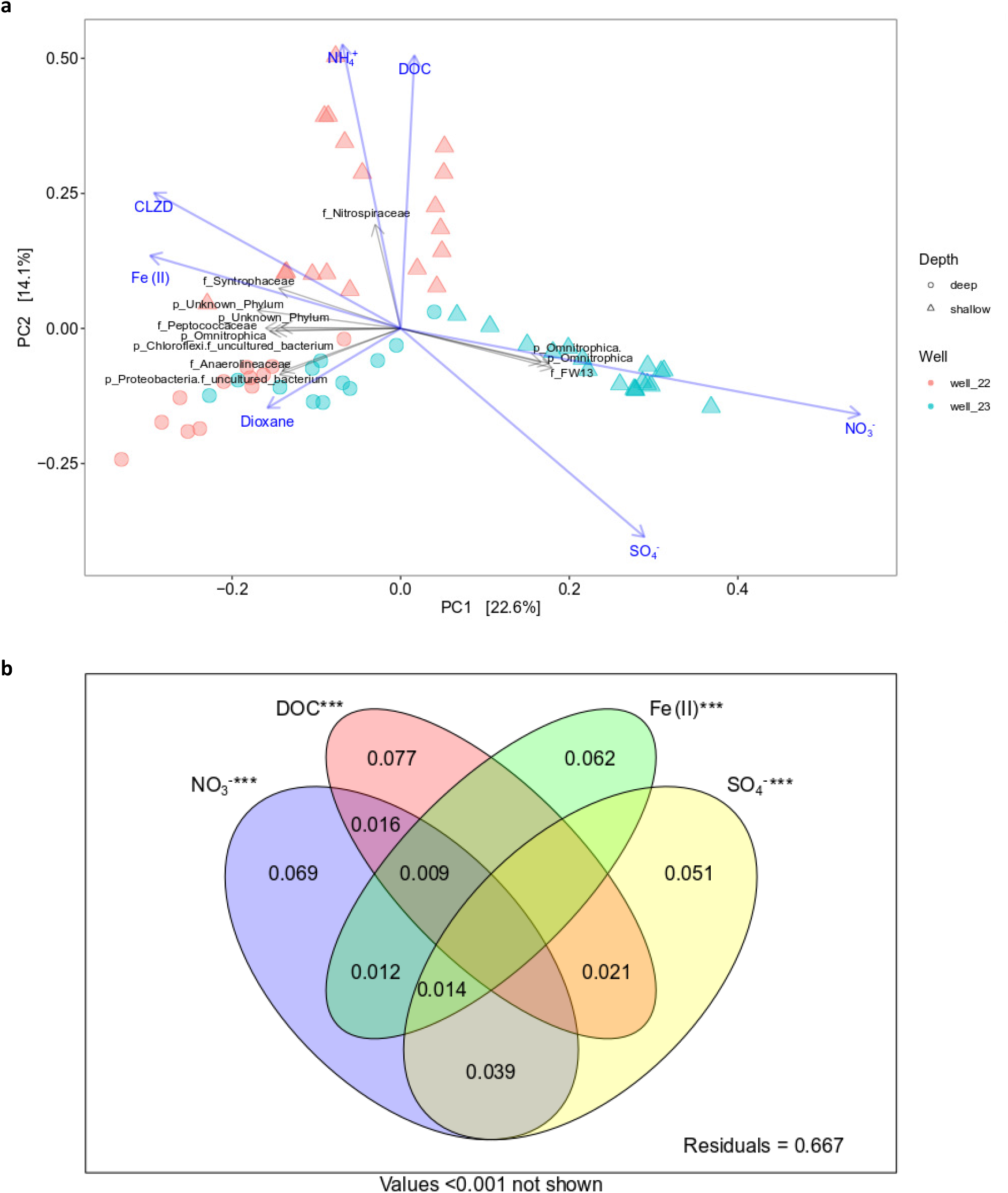
Redundancy analysis of microbial community composition with compounds NO_3_^−^, SO_4_^2−^, Fe(II), DOC, Chloridazon-desphenyl, and 1,4-dioxane as explanatory variables. a. RDA visualizing microbial community composition of samples from all 10 filters (n = 60) colored by well and shaped according to filter (1-3 for shallow, 4-5 for deep). Samples from shallow filters are displayed as triangles, samples from filters as circles. Redundancy analysis (RDA) displays and describes the variation explained in the samples’ microbial communities, given the geochemical compounds and pollutants. Blue arrows indicate geochemical compounds, grey arrows indicate the relative abundance of particular bacterial groups. Length of the arrows is directly proportional to the variance in the microbial community the variable would explain alone. b. Venn diagram visualising the partitioning of the variation and shared variance explained by the significant predictors. *** P<0.001, ** P<0.01, *P<0.05. Values smaller than 0.1% are not shown.

Variation partitioning for nitrate, sulfate, DOC, and iron (II) showed that most of the variation within the microbial communities explained by these compounds was actually only related to one compound (Figure 6b). The biggest shared explained variation was observed for nitrate and sulfate and amounted to 2.2%. Nitrate, sulfate, and DOC shared 1.8% of variation, while the shared variation explained by nitrate, sulfate and iron (II) was 1.7%. The compounds nitrate, DOC, and iron (II) shared 0.6% of the variation, while nitrate and iron (II) shared 0.5% of the variation within the microbial communities.

## 4. Discussion

### 4.1 Selective pressure on microbial communities in aquifers

Groundwater composition was examined to understand microbial community distribution in relation to geochemical parameters. Wells 22 and 23 showed stable geochemical conditions (Figure 1) but varied in pesticide prevalence and concentration over time (Figure 2). In the present study, stable geochemical parameters appear to be the main contributor for stable microbial communities in groundwater ecosystems. Clear and stable redox zonation could be observed with depth (Figure 1), which looks to correlate well with the stable microbial communities observed between 2014-2018 (Figure 4). In contrast, the variation in pesticides presence and concentration over time, unlikely exerted a selective pressure in microbial communities.

This hypothesis was further supported when examining to what extent individual groundwater components could explain the observed variation in microbial community composition. For example, the variation in the microbial community composition was explained mainly by geochemical parameters, namely for 33.3% by DOC, nitrate, sulfate and iron (II) (Figure 6b). In oligotrophic groundwater environments, DOC is an important carbon and energy source supporting metabolic activity (38). In wells studied here, the higher DOC concentrations indicate that DOC was most likely the prevailing carbon substrate, thus limiting microbial use of the MP as carbon source (3). However, DOC can also act as a structural analogue, potentially promoting the development of MP biodegradation capacity (39, 40). The variation in nitrate, sulfate and iron (II) concentrations reflected a redox zonation that appeared to be selective for microbial communities able to utilize the available electron acceptor. This selective pressure was reflected in the qPCR results showing an increase in copy numbers for nitrate or sulfate reduction genes when each of these electron acceptors was present (Figure 1). Even though, qPCR data only show potential catabolic activity, our results confirmed the correlation between geochemical parameters and potential microbial metabolism (3). Thus, in our system, the geochemical conditions appeared to be the main selective driver for shaping microbial community composition.

As opposed to geochemistry, we observed less selective pressure by MP on groundwater microbial community composition. The lack of influence of pesticides on microbial community structure can be explained by the heterogeneous distribution and the low concentrations of these chemicals. Only two of the MPs (CLZD and dioxane) were recurrent enough to allow statistically testing of their influence on microbial composition (Figure 6a). Both MP explained the variation in microbial communities to a lesser extent that geochemical parameters. The explained variation in microbial communities changed from 33.3% (Figure 6b) to 31.6% when sulfate was removed and CLZD was included, and to 32.7% when dioxane but not CLZD was included (Figures S2 and S3). Previous research has shown selective pressure by MP in shallow aerobic aquifers, where microbial communities acclimated to injected MP (41). In that field site, a mixture of pesticides was homogenously distributed by continuous injection for 216 days at concentrations around 40 μg L^−1^ (41), 25 times higher than what we observed in our study. The results of that study are thus in line with ours, namely that chemical compounds present at stable and heightened concentrations exert a selective pressure on microbial communities.

The selective pressure exerted by the availability of electron acceptors is also reflected in key microbial groups (Figure 6a). For instance, we observed two families from the order *Nitrospirales*. *Nitrospiraceae* positively correlated with ammonium presence, and FW13 positively correlated with nitrate presence. These microbial groups are known for participating in different steps of the nitrogen cycle (42, 43). The phylum *Omnitrophica*, previously known as OP3, which is a group of anaerobic microorganisms (44), was also positively correlated to nitrate concentration. The fact that there are microbial groups influenced by nitrate and also a high abundance of nitrate-reducing genes (Figure 1), further confirm the relevance of geochemical conditions for shaping microbial communities and their metabolism. Whether the microbial communities that participate in geochemical cycles have also the potential for MP remediation, needs to be further investigated.

### 4.2 Microbial community distribution with depth in the aquifers

In terms of beta diversity, monitoring wells showed that shallow filters (1–3) in each well cluster separately from deep filters (4–5) from both wells. This pattern was evidenced for samples from 2014-2016 and from 2018, indicating that this finding is not the result of a primer bias. We thus suspect a hydraulic connection between the deeper filters in wells 22 and 23, most likely caused by the large volumes of water extracted in the deeper aquifer. The possibility of a hydrological connection implies that MP found in the deeper filters can spread more easily through the aquifer.

To corroborate whether there is a possible hydrological connection between the two studied wells, the soil profile from both wells was obtained from a project partner (Tables S16 and S17). Both wells were mainly composed by sand, but there were also clay layers dividing shallow (1–3) from deep (4–5) filters. Compressed clay layers can be enclosing part of the aquifer (45) and therefore potentially influencing the differences in microbial composition. Furthermore, in the deeper section (40-60 m), the sand was coarse, meaning that groundwater could easily flow throw the soil particles. The implication of this hydrological connection for the application of remediation technologies needs to be considered.

### 4.3 Natural attenuation future perspectives

Exploring the microbial communities is the first step to understanding the MP biodegradation potential in aquifers. The application of monitored natural attenuation as a remediation technology, relies mainly on tools that are still not fully developed for groundwater systems. For demonstrating MP natural attenuation, there are three lines of evidence suggested: i) changes in MP concentration or biodegradation products, ii) changes in geochemical data as indirect evidence, and iii) *in situ* or microcosm experiments that provide direct evidence of biodegradation (46). Measuring changes of MP concentrations in groundwater systems may not provide evidence of biodegradation, since MP can be heterogeneously distributed, as observed in our dataset (Figure 1). Degradation products are also difficult to identify since anaerobic degradation pathways are largely unknown (47). We showed that geochemical data can be stable over time (Figure 1). However, if biodegradation happens, it is unlikely that it results in geochemistry changes. First, because concentrations of MP are generally three orders of magnitude lower than those of geochemical parameters and second because in the presence of DOC, MP will not be consumed at high rates. Even when microcosm experiments can provide direct evidence for degradation, this can mainly be demonstrated by changes in MP concentration and changes in known degradation gene abundance (38). In the field, the systems are more complex, therefore those two tools will not be enough to prove biodegradation.

Remediation technologies such as biostimulation and bioaugmentation are also an option to remove MP from groundwater systems. In the present dataset, there are a variety of redox conditions for biodegradation to occur. Still, there is lack of information about which redox condition could facilitate degradation of a specific pesticide. In principle, understanding the right conditions for biodegradation would allow us to engineer the system. By the use of biostimulation, for instance, the right redox can be provided for the microbial communities depending on which pesticide is present. In this particular example, we could monitor changes in the MP concentration per depth and changes in the stable geochemical data. If we incorporate the information we have about the horizontal hydrological connection in the deeper aquifer, we could determine at which depth it could be more effective to provide the amendment. Furthermore, the incorporation of additional monitoring tools, such as compound specific isotope analysis (CSIA) (48, 49) can provide a stronger argument about whether engineering the system was or not successful.

### 4.4 Outlook and recommendations

The aim of this study was to explore groundwater microbial communities for natural attenuation potential of MP. The present study provides information about the main microbial groups present in two groundwater monitoring wells, and the influence of geochemical parameters and MP in the microbial community composition. We found that the main selective pressure in the aquifer were the geochemical conditions. MP were heterogeneously distributed and at low concentrations, thus showing less effect on the microbial composition. The results shown here can be used for the design of degradation experiments of MP under different redox conditions as well as for inspiration to use molecular tools for monitored natural attenuation projects.

Our results confirmed that microbial groups in groundwater have yet to be sufficiently explored, as exemplified in the high abundance of sequences that could not be classified at the phylum level (Figure 3). The lack of anaerobic MP degraders described in literature, makes the use of molecular targeted tools difficult since many degraders have not been described before or they might be clustered together in “unknown” taxa. Application of high throughput and non-targeted technologies such as metagenomics, can contribute to explore the genetic degradation potential in groundwater systems. We suggest for future studies of unexplored environments the combination of targeted (i.e., 16s rRNA, qPCR) and untargeted molecular tools (i.e., metagenomics) to determine the presence of known microorganisms and the metabolic capacity of both known and unknown. Within our dataset, we tried to understand the effect of certain MP on microbial communities. This approach is thus limited to the MP included in this dataset and cannot discard the possibility of unidentified MP or degradation products to have a different effect in the microbial community structure. It is suggested that a bigger set of emerging MP is included in future research projects. This study provides initial insights on how the microbial communities are composed and distributed in groundwater systems and what are the main factors influencing them. We aim in the future to conduct further studies towards understanding the natural attenuation capacity of groundwater in order to guarantee safe drinking water production from aquifers.

## 5. Acknowledgments

This research is funded by TKI project MicroNAC and NWO Veni grant 15120. We would like to acknowledge drinking water institutions Water Laboratorium Noord and Vitens for providing support of this project, including groundwater monitoring data and logistical support of field sampling campaigns. Special thanks to Albert-Jan Roelofs (WLN, The Netherlands) for his help during all field work. We also thank Laura Piai, Koen van Gijn and Thomas Wagner (Wageningen University, The Netherlands) for helping us improve the manuscript quality.

